# Collagen crosslinking and organizational patterns reflect common disease processes in idiopathic pulmonary fibrosis and non-resolving acute respiratory distress syndrome

**DOI:** 10.64898/2026.05.09.723675

**Authors:** M. Nizamoglu, O. A. Carpaij, T. Borghuis, Judith M. Vonk, R. M. Jongman, M. C. Morrison, R. Hanemaaijer, P. J. Wolters, J. Pillay, J.K. Burgess

**Author notes:** **Corresponding author: Janesh Pillay**, Department of Critical Care, University Medical Center Groningen, Hanzeplein 1, 9713 GZ, Groningen, the Netherlands. Contributed equally (J. Pillay, J.K. Burgess).

## Abstract

**Rationale:** Fibrotic lung diseases, such as idiopathic pulmonary fibrosis (IPF) and fibroproliferative remodeling in acute respiratory distress syndrome (ARDS), are characterized by increased extracellular matrix (ECM) deposition. However, measuring collagen accumulation alone does not capture differences in ECM organization or biochemical maturation that may distinguish persistent fibrosis from potentially reversible remodeling.

**Objectives:** To examine collagen organization characteristics and mature (pyridinoline) collagen crosslinking amount in established end stage fibrotic lung disease (IPF) and fibroproliferation following an acutely damaged lung (non-resolving (NR) ARDS) and to investigate any relationships in these parameters and temporal tissue remodeling.

**Methods:** Human lung tissue samples from control subjects, patients with IPF, and NR-ARDS were analyzed. Collagen amount and fiber organization were digitally quantified using picrosirius red staining. Mature collagen crosslinking was assessed by quantification of pyridinoline crosslinks.

**Measurements and Main Results:** Lung tissue from both IPF and NR-ARDS lungs had higher collagen content compared with controls. Collagen fiber organization differed between groups. IPF lungs exhibited collagen architectures consistent with established fibrosis, whereas NR-ARDS lungs showed altered but less stabilized collagen organization despite similarly elevated collagen levels. Mature collagen crosslinks were significantly higher in IPF lungs but not in NR-ARDS lungs compared to controls. Integrated analyses identified distinct disease-associated ECM phenotypes, indicating that higher collagen abundance in NR-ARDS, unlike IPF, is not accompanied by more mature and persistent collagen crosslinking.

**Conclusions:** Despite shared increases in collagen content, IPF and NR-ARDS lungs differ fundamentally in collagen organization and crosslinking maturity, suggesting differences in the reversibility of these conditions.

## Introduction

Tissue remodeling in the lung is a core process to maintain homeostasis and pulmonary integrity. Disruption of this process represents common pathways shared across many lung diseases, including idiopathic pulmonary fibrosis (IPF) and fibroproliferative remodeling in acute respiratory distress syndrome (ARDS) (1-3), even when the initiating triggers may differ (4, 5).

In IPF this unbalanced tissue remodeling drives progressive lung function decline and a median survival of 2.9 – 4.4 years after diagnosis, making this prognosis worse than most cancers (6). There is no curative medical therapy for IPF beyond lung transplantation (7). Similar tissue remodeling can be observed in persistent or non-resolving ARDS (NR-ARDS) which becomes more apparent in prolonged disease courses (3, 8). Excessive tissue remodeling in ARDS is associated with increased mortality and functional impairments in survivors (9). To what extent this tissue remodeling in NR-ARDS resembles irreversible fibrosis seen in IPF is unknown. Implied irreversibility in NR-ARDS, based on HRCT criteria of fibrosis (e.g. reticulation, traction bronchiectasis), can prompt limitation or withdrawal of supportive care, or lung transplantation in selected centers.

Historically, fibrosis is quantified through measuring collagen accumulation using histologic analysis or biochemical and analytical approaches (10, 11). This is often performed in tandem with measurements associated with collagen synthesis, including post-translational modification processes or collagen turnover fragments released during matrix remodeling (12-15). Recent studies expose the importance of considering organizational properties of the ECM, as collagen amount alone reveals only part of the complex fibrotic landscape (5, 16). This distinction is important, as organizational structure of the collagen network can promote further fibrotic responses in resident and infiltrating cells (17-20). However, the relationship between the amount and the organization of lung collagens during fibrotic remodeling remains under-explored.

Collagen organization is impacted by several factors. Reversible, non-covalent bonds such as hydrogen bonds or ionic interactions are key factors in collagen assembly (21, 22). ECM crosslinking enzymes, including lysyl oxidase (LOX) and LOX-like family members and transglutaminases, link ECM molecules, including collagens, to each other through covalent bonding at defined amino acids (23-26). Over time, irreversible mature pyridinoline crosslinks form and accumulate, providing an indication of prolonged collagen remodeling (27, 28). While changes in collagen organization during lung diseases are recognized (29), thorough characterization of lung tissue regarding collagen organization and mature collagen crosslinking remains limited.

In this study, we hypothesized that the pyridinoline crosslinking in lung pathologies differs among control, NR-ARDS, and IPF lung tissue, and adds information beyond the high collagen amount observed in IPF and NR-ARDS compared to control lungs. To explore this, we measured the collagen amount, collagen fiber assembly, and mature collagen crosslinking in lung tissue samples from these groups. Through coupling the collagen organization parameters to the amount of mature collagen crosslinking, we provide insights on disease-associated ECM phenotypes that extend beyond collagen amount and may refine how fibrotic lung tissue remodeling is interpreted.

## Methods

### 1. Lung tissue collection, processing, and experimental design

Control lung samples were obtained from macroscopically normal regions from oncological lung resections. IPF lung samples were sourced from explanted lungs during transplantation surgeries. SARS-COV2 induced (COVID)-ARDS lung tissue samples were obtained from explanted lungs, partly provided by the UCSF University of California. Non-COVID ARDS lung samples were from explanted lungs obtained post-transplant or post-mortem. This study was conducted in compliance with the UMCG Research Code (https://umcgresearch.org/w/research-code-umcg) and the Dutch Code of Conduct for Health Research (https://www.coreon.org/wp-content/uploads/2023/06/Code-of-Conduct-for-Health-Research-2022.pdf). Use of clinical data and leftover lung tissue was exempt from the Dutch Medical Research Involving Human Subjects Act (WMO), as confirmed by the UMCG Medical Ethics Committee. The protocols were approved by the UMCG Central Ethics Review Board for non-WMO studies (study number 10748 and 18260) and, under Dutch law (WGBO art. 458; GDPR art. 9; UAVG art. 24), did not require informed consent. The protocol concerning the samples from the University of California, San Francisco (UCSF) was approved by the Institutional Review Board of USCF (approval number: 13-10738). All donor material and clinical data were deidentified prior to experimental use. An overview of patient characteristics is provided in **Table 1**. Lung tissue samples were fixed in formalin and subsequently embedded in paraffin, in line with the protocols described previously (30). The experimental approach is outlined in **Figure 1**.

**Table 1:**
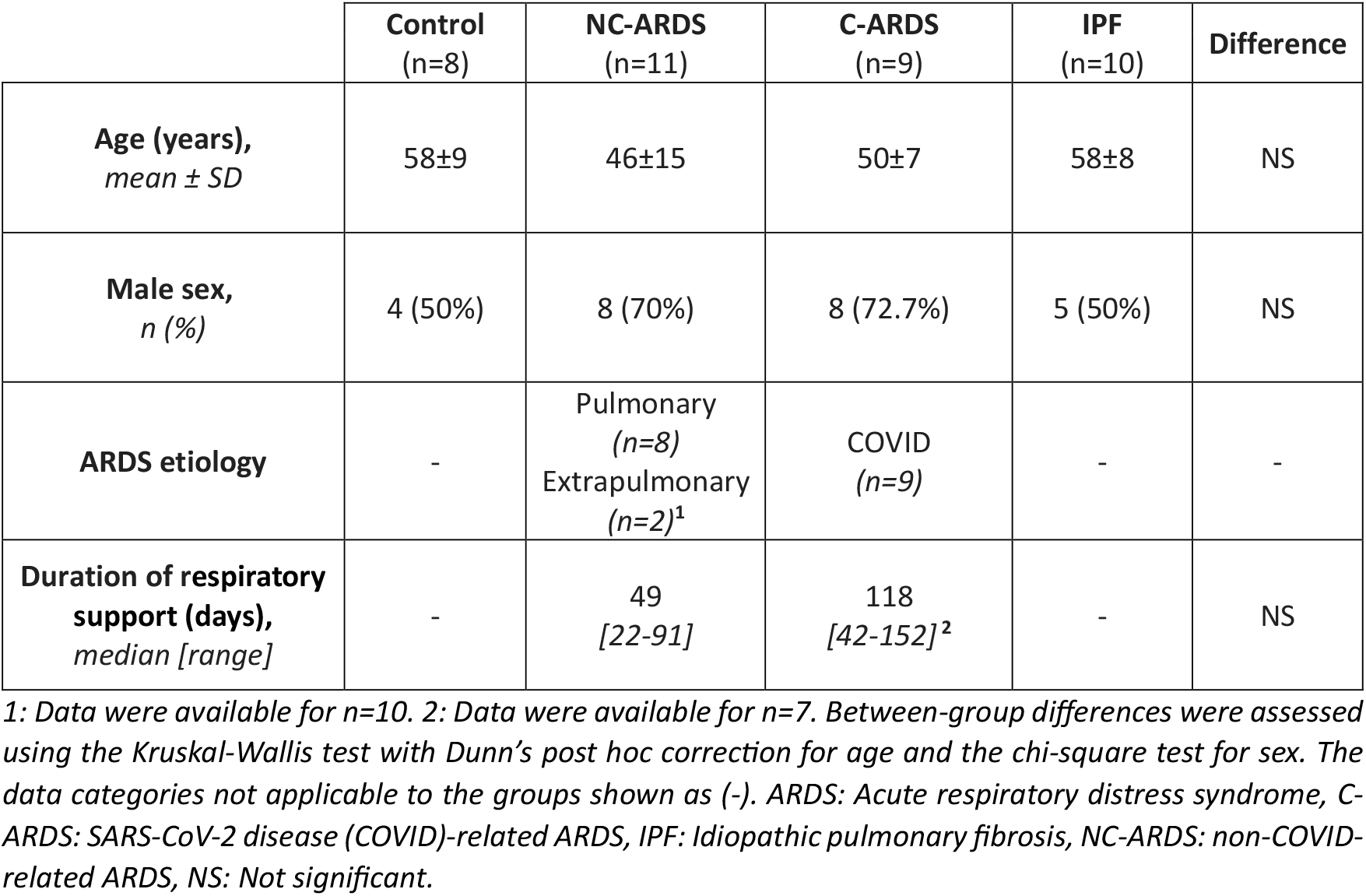
Patient characteristics of tissue donors of the samples used in this study.

**Figure 1:**
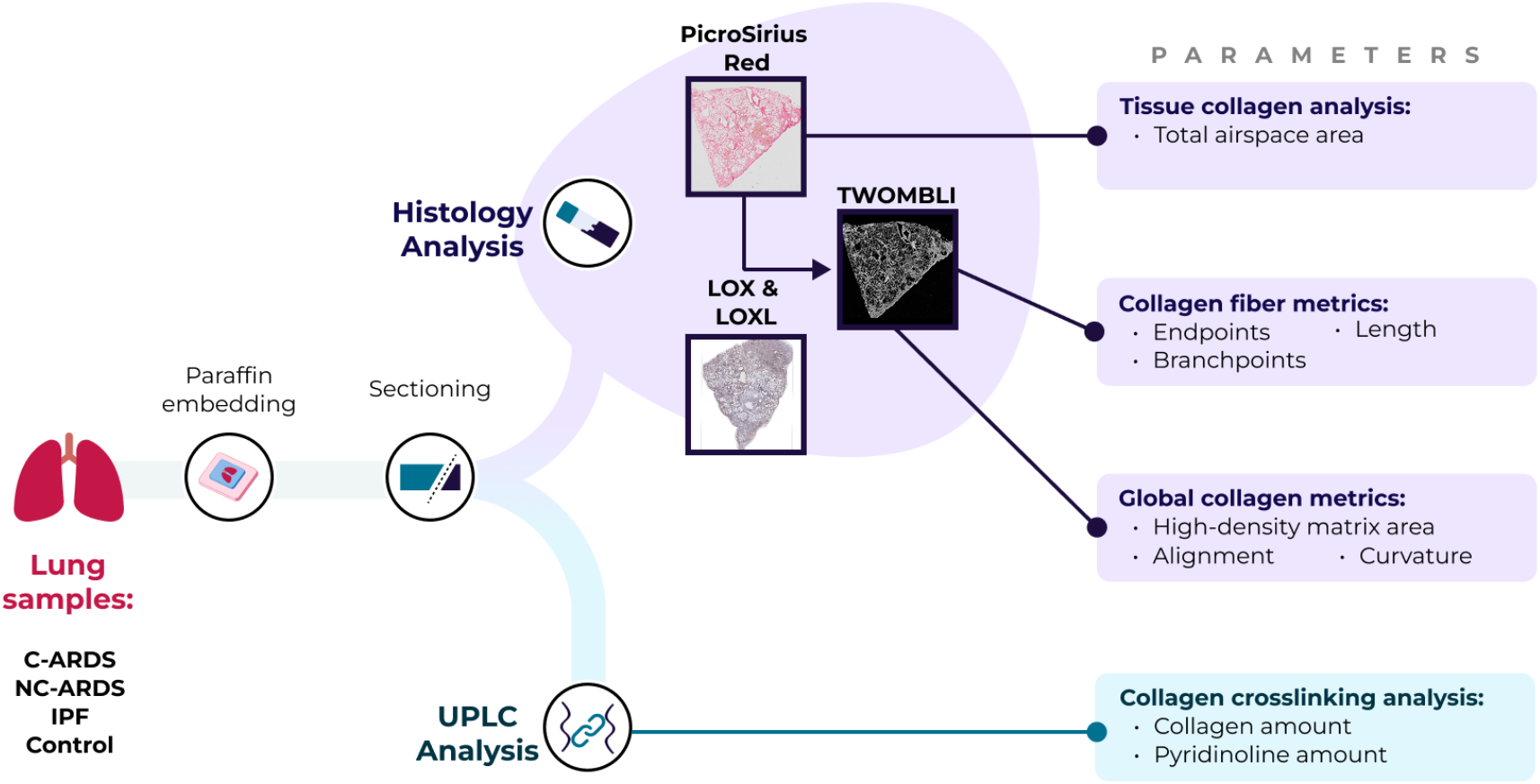
Experimental design and parameters investigated in this study. Lung samples from control, NC-ARDS, C-ARDS, and IPF were paraffin-embedded and sectioned for histological and biochemical analyses. Sections stained with Picrosirius Red were used to assess total collagen, as well as the airspace through the amount of free space. Collagen fiber architecture was quantified using TWOMBLI, extracting fiber-level metrics (endpoints, branchpoints, length) and global matrix features (high-density matrix area, alignment, curvature). IHC staining for LOX and LOXL members were performed on serial sections. In parallel, UPLC analysis quantified total collagen and pyridinoline collagen crosslinking. ARDS: Acute respiratory distress syndrome, C-ARDS: SARS-CoV-2 disease (COVID)-related ARDS, IHC: Immunohistochemistry, IPF: Idiopathic pulmonary fibrosis, LOX: Lysyl oxidase, LOXL: LOX-like, NC-ARDS: non-COVID-related ARDS, UPLC: Ultra-performance liquid chromatography TWOMBLI: The Workflow Of Matrix BioLogy Informatics [from (35)].

### 2. Immunohistochemical staining

Five μm sections of lung samples were deparaffinized. Hematoxylin and eosin (H&E) staining for tissue visualization and staining using 0.1% PicroSirius Red solution (PSR; Sigma St. Louis, MO, USA) for total collagen visualization were performed as described elsewhere (19). Staining for lysyl oxidase (LOX), LOX-like 1 (LOXL-1), and LOX-like 2 (LOXL2) was performed following procedures described elsewhere (31, 32). For primary antibodies, 1:500 rabbit anti-LOX (catnr: ab31238, Abcam, Cambridge, UK), 1:800 rabbit anti-LOXL1 (catnr: PA5-79609, Invitrogen, California, USA), and 1:500 rabbit anti-LOXL2 (catnr: NBP1-32954, Novus Biologicals, Centennial, CO, USA) dilutions were used. Positive staining on target proteins was visualized using Vector NovaRED (catnr: SK-4800, Vector Laboratories, Newark, USA). Tissue sections were imaged using a Hamamatsu NanoZoomer digital scanner (Hamamatsu Photonic K.K., Shizuoka, Japan) (33). Fluorescence imaging of PSR-stained sections was performed using Olympus VS200 slide scanner (Hamburg, Germany).

### 3. Collagen and collagen crosslinking measurement

Quantification of total collagen and pyridinoline crosslinking was performed through ultra-performance liquid chromatography (UPLC) as previously described (34). Briefly, three serial 15 μm sections from paraffin-embedded lung samples (a total of 45 μm tissue depth) per donor were cut, deparaffinized, homogenized and then digested in 6M HCl. Total collagen amount per sample was measured using a sensitive tissue collagen assay (Quickzyme, Leiden, the Netherlands) following the manufacturer’s instructions. Then the samples were dried under nitrogen and reconstituted in elution buffer (48007, Chromsystems Instruments & Chemicals GmbH, Gräfelfing, Germany). Pyridinoline crosslinks were quantified by UPLC (ACQUITY, Waters Chromatography B.V., Breda, the Netherlands) using a high-performance column (48100, Chromsystems) with isocratic separation at a 1.2 mL/min flow rate (using mobile phase 48001, Chromsystems). Detection of pyridinoline crosslinks was performed with a fluorescence detector (ACQUITY UPLC Fluorescence Detector, Waters Chromatography) set at 290 nm excitation and 395 nm emission. Mature crosslinks were expressed as mol crosslinks per mol collagen (mol/mol).

### 4. Image analysis

Digital image analysis of the light microscopy images was performed using an in-house macro using Fiji Image J (v1.53F51) (36). Airspace area was calculated by measuring the area inside the manually selected exterior boundary of the total tissue region (including empty space) and subtracting the measured tissue-positive area. The fluorescence images of PSR-stained tissue sections were first divided to 1500 x 1500-pixel tiles using QuPath (v0.5.1). Tiles were then analyzed using The Workflow Of Matrix BioLogy Informatics (TWOMBLI) plugin in Fiji ImageJ following developers’ instructions as described elsewhere (35). Tiles that were technically unsound and not able to be processed by TWOMBLI (1.2% of all tiles) were deleted from the analysis batch. All images were analyzed using the standard parametric thresholds.

### 5. Statistical analyses

Statistical analyses of the data generated were performed using an interaction analysis in a linear mixed model analysis in IBM SPSS Statistics (v30.0.0; IBM, Armonk, New York, USA) to account for multiple tiles per section belonging to one tissue sample. Normality of the data was tested through Q-Q plots and Shapiro-Wilk test. TWOMBLI results, airspace area, and pyridinoline crosslinking data were log-transformed before analyses. All results are presented as mean ± standard deviation. Missing donor-level feature values were imputed using feature-wise medians before z-score normalization. PCA was then performed on the standardized donor-by-feature matrix as an exploratory dimensionality-reduction method to project correlated quantitative features into orthogonal principal components. This was done to visualize the dominant axes of multivariate variance across donors without imposing group labels. In total, 36 of 380 donor-by-feature values were imputed (9.47%). Plotting of the graphs and subsequent correlations and PCA were done using Python (v3.13.1) and the following packages: NumPy (v2.1.3), Pandas (v2.2.3), Matplotlib (3.10.0), scikit-learn (v1.6.1), and SciPy (v1.15.1).

## Results

### 1) Collagen amount is higher in both ARDS and IPF lung samples

To visualize the collagen content and distribution within the lung samples, we used a PicroSirius Red staining, identifying all collagens (**Figure 2A-D**). Control lung sections revealed a physiologically normal distribution of collagen: thin and delicate presence around the alveoli and thicker layers around the airways and blood vessels. The images of the sections of IPF lung tissues showed deposition of aberrant ECM in the interstitial regions. For the ARDS groups, we observed accumulation of collagen in the interstitial regions in both NC-ARDS and C-ARDS samples. We then quantified the total collagen amounts in these tissue sections (**Figure 2E**). Among the four groups, IPF lung samples had the highest collagen content (1910.08 ± 520.33 µg/mL), significantly exceeding that of control lung samples (417.12 ± 294.88 µg/mL; p<0.001; linear mixed model analysis), which had the lowest collagen levels. Both NC-ARDS and C-ARDS sections had higher levels of collagen compared to control lung sections (p<0.001, and p=0.005, respectively). The levels of collagen did not differ between the ARDS groups, but these values were lower than those in IPF (p=0.002 and p<0.001, respectively).

**Figure 2:**
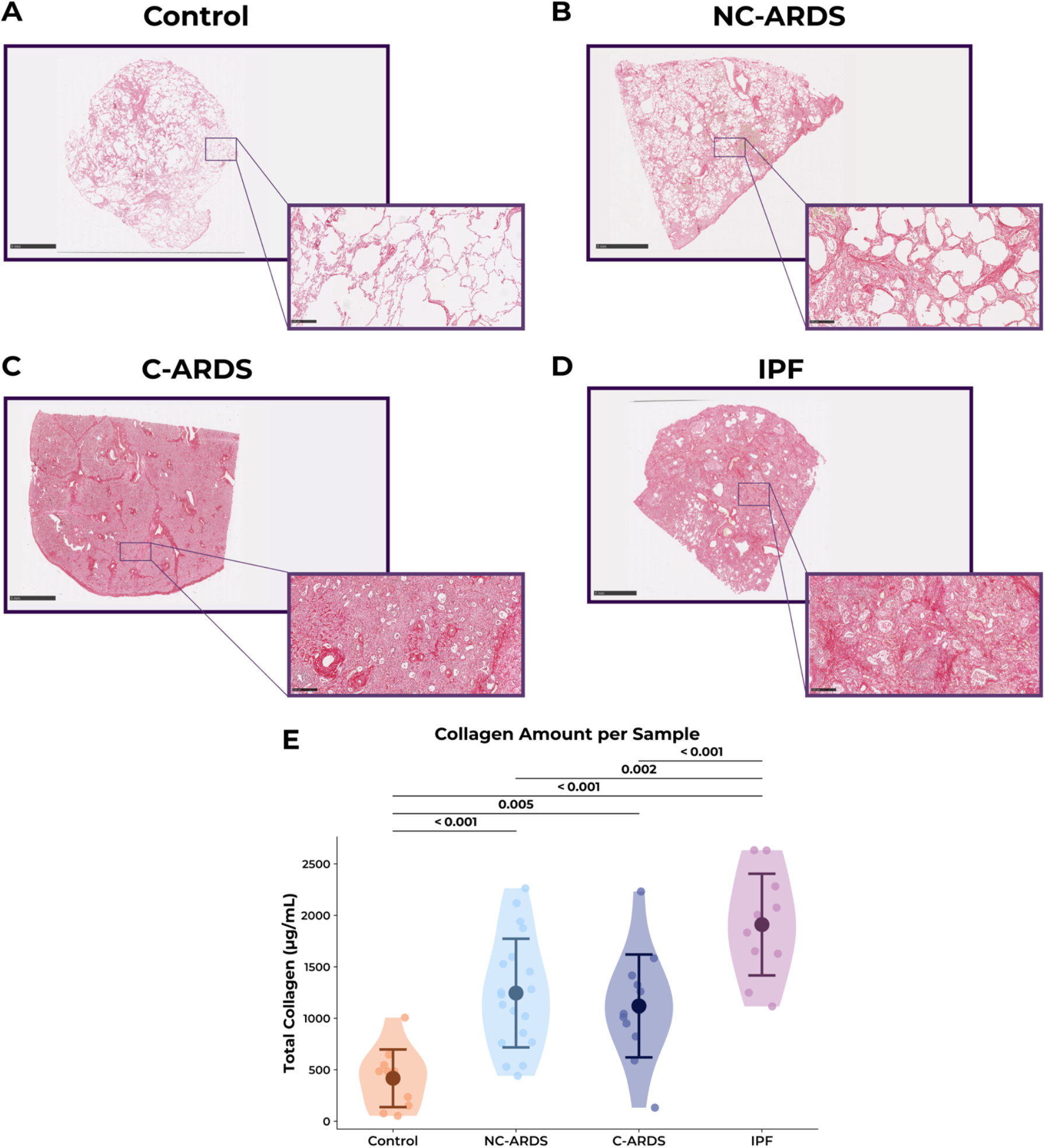
Higher collagen amount in both IPF and ARDS lungs samples, than controls. Example images of A) control, B) non-COVID ARDS, C) COVID ARDS, and D) IPF lung tissue sections stained with PicroSirius Red to visualize collagens. Scale bars=5 mm for zoomed out panels and 250 μm for zoomed in panels. E) Quantification of total collagen content through biochemical analysis. Data represent individual measurements from 1-2 samples per patient. Group mean ± SD values are superimposed on the violin plots. Sample sizes were n=10 for control samples, n=19 for non-COVID ARDS, n=12 for COVID ARDS, and n=10 for IPF. These samples were obtained from 7 individuals in the control group, 11 patients with non-COVID ARDS, 9 patients with COVID ARDS, and 7 patients with IPF. A mixed model analysis was used to assess the statistical significances while accounting the variance from multiple sections per patient. ARDS: Acute respiratory distress syndrome, C-ARDS: SARS-CoV-2 disease (COVID)-related ARDS, IPF: Idiopathic pulmonary fibrosis, NC-ARDS: non-COVID-related ARDS.

### 2) Collagen organization patterns were different across the lung diseases

Next, we examined if ECM organization differed between control, ARDS or IPF lungs. Starting with airspace area (**Figure 3A**), we observed an inverse pattern compared to total collagen amount, where the control lungs had the greatest percentage airspace (65.95% ± 21.95). All three diseased lung sample groups had a lower percentage airspace than the controls (p=0.004 for NC-ARDS, p<0.001 for C-ARDS, and p=0.003 for IPF).

**Figure 3:**
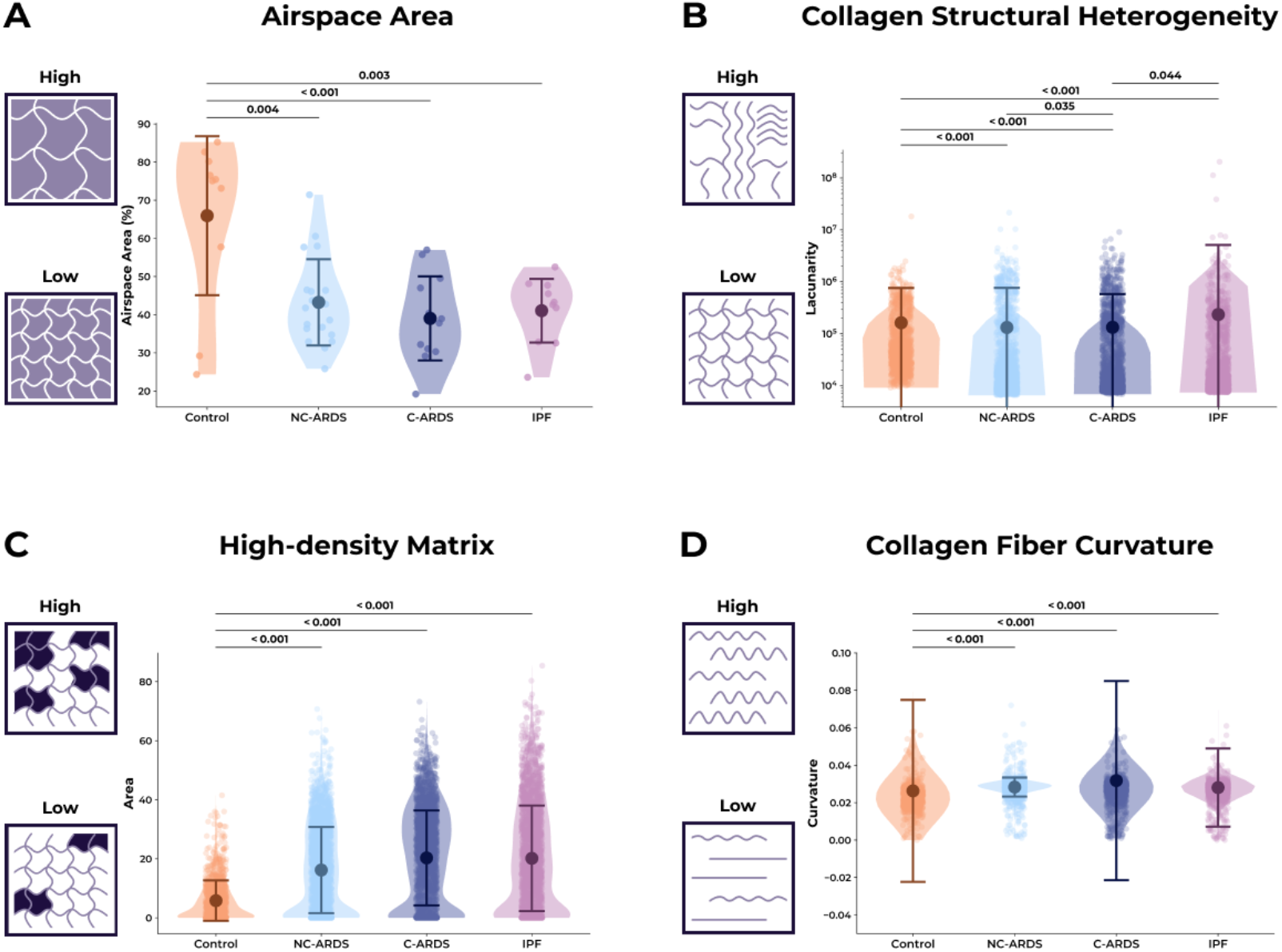
Collagen organization patterns differ in ARDS and IPF lung tissues. The collagen organization analysis was performed through TWOMBLI analysis on images of PSR-stained lung sections. Schematic explanation and quantification of A) airspace area, B) collagen distribution heterogeneity, C) high-density collagen matrix areas, D) collagen fiber curvature. Data represent individual tiles from 1-2 images per patient. Group mean ± SD values are superimposed on the violin plots. For panel A, sample numbers were 10 for control, 19 for non-COVID ARDS, 12 for COVID ARDS, and 10 for IPF groups, derived from 8, 10, 9, and 10 individuals, respectively. For panels B to D, the analysis included 6 control individuals, 10 patients with NC-ARDS, 9 patients with C-ARDS, and 9 patients with IPF. Total image tiles analyzed, with per-individual ranges in parentheses, were as follows: control, 1,087 (90 to 415); NC-ARDS, 2,406 (132 to 405); C-ARDS, 2,374 (102 to 552); and IPF, 2,349 (101 to 372). A mixed model analysis was used to assess the statistical significances while accounting for the variance from multiple tiles and images per sample. ARDS: Acute respiratory distress syndrome, C-ARDS: SARS-CoV-2 disease (COVID)-related ARDS, IPF: Idiopathic pulmonary fibrosis, NC-ARDS: non-COVID-related ARDS.

Collagen assembly patterns were compared between control and diseased lungs using automated image analysis of PSR-stained sections. The heterogeneity of the structural arrangement of the collagen fibers was measured as the patchiness of collagen distribution (lacunarity) in the tissue images (**Figure 3B**). Control lungs had the lowest collagen structural heterogeneity compared to other groups (p<0.001 for all groups). C-ARDS lungs had less collagen structural heterogeneity than both NC-ARDS lungs (p=0.035), and IPF lungs (p=0.044). The next parameter we checked was high-density matrix, quantifying the areas of densely packed collagen. Control lungs had the lowest proportion of high-density matrix compared to the three disease groups (p<0.001 for all pairwise comparisons; **Figure 3C**). This was consistent with a denser overall collagen fiber network in diseased lungs (*Supplementary Figure 1A*). Characterizing the ECM organization through the collagen fiber curvature (**Figure 3D**) revealed that control lungs had a lower fiber curvature than the three diseased groups. This trend was also present in the number of collagen fiber branchpoints (*Supplementary Figure 1B*) and endpoints (*Supplementary Figure 1C*), but not in collagen fiber alignment (*Supplementary Figure 1D*). These findings collectively show that collagen assembly differs between ARDS and IPF despite elevated collagen levels in both diseases.

### 3) IPF lungs had the highest mature crosslinking levels

We examined the presence of LOX, LOXL1 and LOXL2 in sections of lung tissue samples as potential crosslinking enzymes. LOX staining was variable across fibrotic lung tissue samples, with epithelial staining frequently observed but not consistently present across all samples (*Supplementary Figure 2*). In ARDS lung tissue sections, LOX staining appeared relatively low. LOXL1- and LOXL2-stained sections showed heterogeneous presence and distribution among all lung tissue sample types (*Supplementary Figures 3 & 4*, respectively).

Mature collagen crosslinking in lung samples was assessed by quantifying pyridinoline crosslinks, a biochemical marker of collagen maturation (**Figure 4A**). We quantified the mature collagen crosslinks per collagen molecule using UPLC (**Figure 4B)**. IPF lungs had the highest levels of mature crosslinks (0.357 ± 0.08 mol/mol) compared to control (p=0.005), NC-ARDS (p<0.001), and C-ARDS (p=0.017) lungs. Overall, crosslinking per collagen levels in both NC-ARDS and C-ARDS lungs were similar to the control lungs (0.23 ± 0.04 mol/mol) and did not differ from each other (0.24 ± 0.11 mol/mol and 0.26 ± 0.08 mol/mol, respectively). In NR-ARDS, despite heterogeneity in the data, there was a relationship between time in ICU and total collagen amount in the lung tissue (*Supplementary Figure 5A*). Once divided into three time-banded subgroups, there was a trend between the mature collagen crosslinks and ICU duration (*Supplementary Figure 5B*) When the relationship between the collagen amount and the crosslinking amount was investigated, in IPF there was a positive relationship between the amount of collagen and the degree of crosslinking per collagen molecule. In NR-ARDS the opposite pattern was present, with a lower level of crosslinking per molecule when more collagen was present (**Figure 4C**).

**Figure 4:**
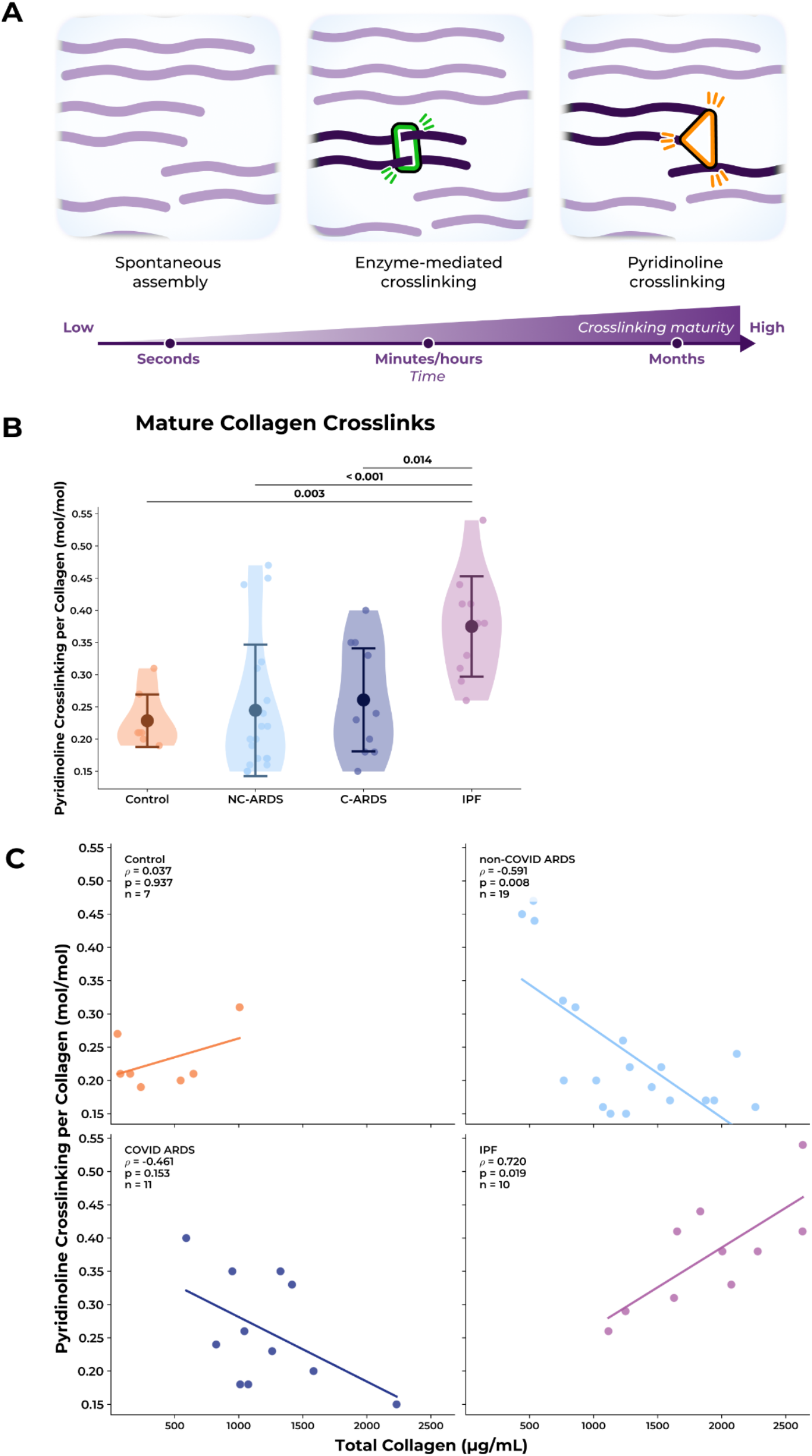
Mature collagen crosslinking is altered in ARDS, compared to established fibrosis in IPF and to control lungs. A) Schematic representation of premature and mature collagen crosslinking. B) Quantification of mature collagen crosslinking through pyridinoline crosslinks per collagen molecule in the lung sections. Data represent individual measurements from 1-2 samples per patient. Group mean ± SD values are superimposed on the violin plots. Sample sizes were n=7 for control, n=19 for non-COVID ARDS, n=11 for COVID ARDS, and n=10 for IPF, derived from 6 control individuals, 10 patients with non-COVID ARDS, 8 patients with COVID ARDS, and 10 patients with IPF. C) Association between total collagen present in lung tissue section and crosslinks per collagen molecule across the disease groups A mixed model analysis was used to assess the statistical significances while accounting the variance from multiple sections per patient. ARDS: Acute respiratory distress syndrome, C-ARDS: SARS-CoV-2 disease (COVID)-related ARDS, IPF: Idiopathic pulmonary fibrosis, NC-ARDS: non-COVID-related ARDS.

**Figure 5:**
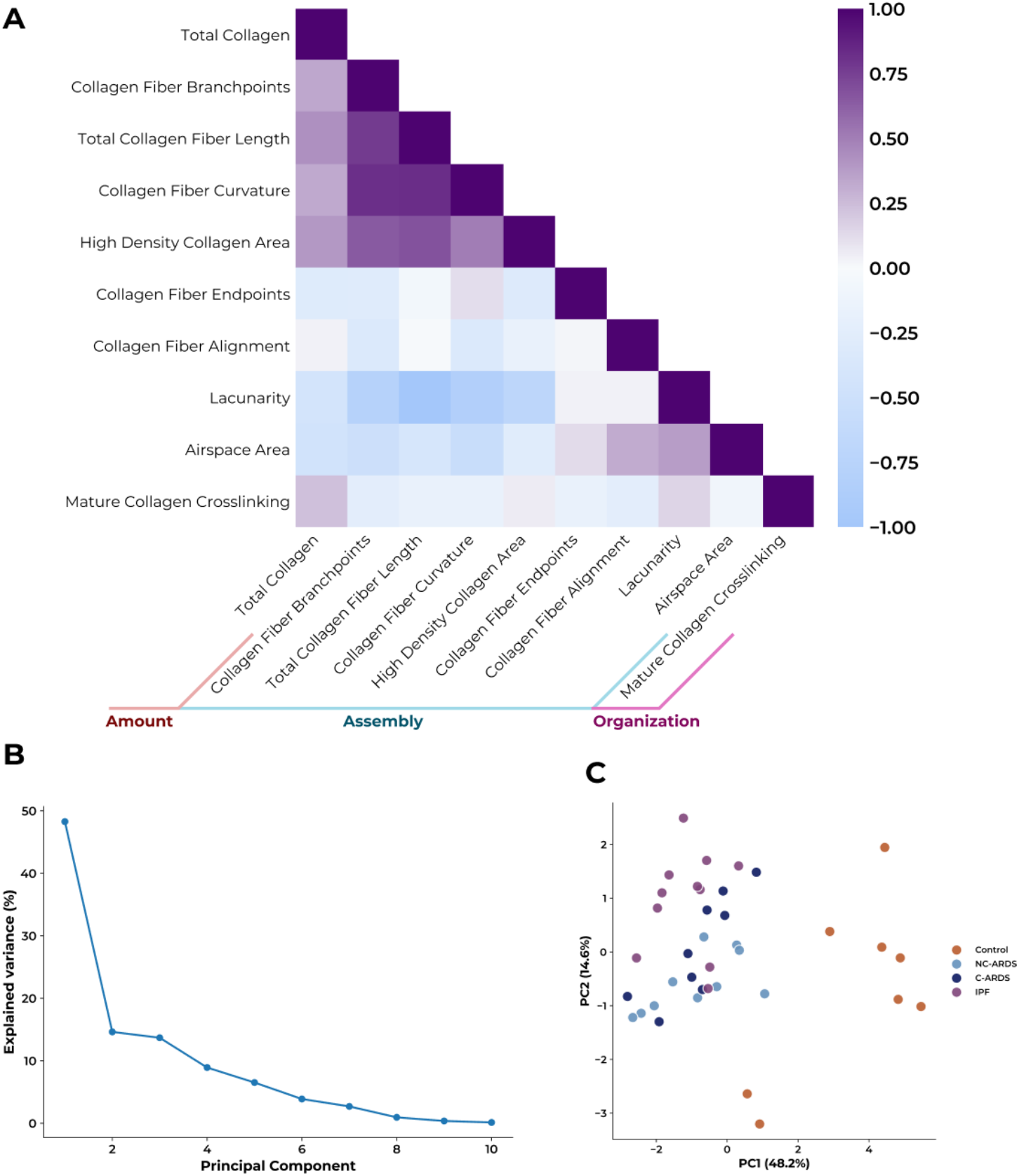
Collagen architecture metrics reveal distinct ECM remodeling patterns across disease groups. A) Correlation heatmap describing the co-occurrence of different parameters belonging to collagen amount, assembly, and organization categories measured in this study. Only the lower triangle is shown for clarity. B) Scree plot showing the proportion of variance explained by the first 10 principal components. Principal component 1 (PC1) explains 48.2% of the total variance and principal component 2 (PC2) explains 14.6%. B) Scores plot of PC1 versus PC2 demonstrating separation among control, NC-ARDS, C-ARDS, and IPF lung samples. Each point represents an individual donor (n=7 control, n=10 non-COVID ARDS, n=9 COVID ARDS, n=10 IPF). ARDS: Acute respiratory distress syndrome, C-ARDS: SARS-CoV-2 disease (COVID)-related ARDS, IPF: Idiopathic pulmonary fibrosis, NC-ARDS: non-COVID-related ARDS.

### 4) Interplay between Collagen Crosslinking and Organization

To examine the interplay between the mature collagen crosslinking and the collagen assembly parameters we quantified in this study, we first checked which parameters were correlating with each other (**Figure 5A**; correlation coefficients and associated statistical tests are reported in *Supplementary Figure 6*). Higher total collagen amount was associated with a denser and more complex collagen network, reflected by greater fiber length, branching, curvature, and high-density collagen area, and by lower lacunarity and airspace area. Most fiber-based parameters were positively correlated with each other, whereas fiber alignment showed weak associations with the remaining metrics. Lacunarity and airspace area were inversely related to most fiber-based parameters, consistent with a more open matrix architecture. Mature collagen crosslinking showed weak associations with the structural parameters and no clear relationship with total collagen amount.

In order to further understand the co-occurrence of the collagen assembly parameters and the crosslinking characteristics, we performed a principal component analysis (PCA): principal component 1 (PC1) accounted for the 48.2% of the total variance while PC2 explained 14.6% of the variance in the samples (**Figure 5B**). Patient samples also showed a pathology-driven clustering based on these identified PCs (**Figure 5C**). Together, these illustrate the patterns that represent the differences that occur in the coordinated collagen remodeling among fibrotic lungs due to different pathologies and temporal dynamic differences in the disease courses.

## Discussion

In this study, we investigated ECM organization and crosslinking in control, NR-ARDS and IPF lung samples through digital image analysis. We showed that while there was an overall increase in collagen amount in the diseased lung samples, there were distinct structural organizational changes that differed between the diseases. Increased collagen content was observed in NR-ARDS lungs which showed a remodeled tissue structure in which the excess collagen fibers had not developed mature chemical crosslinks. IPF lungs showed the most advanced fibrotic phenotype, with very high collagen content, marked architectural distortion, and the highest level of mature collagen crosslinking. Correlations were found between different collagen structural parameters which translated to distinct patterns in the tissue structural changes that characterized the altered collagen matrix in NR-ARDS and IPF. These findings indicate there are defined patterns of collagen assembly and organization in fibrotic lung diseases that are reflective of the maturity of the collagen crosslinking, expanding our understanding of the collagen network structure in fibrotic responses in the lung and the potential for reversing pathologic collagen deposition.

Increased collagen was present in diseased lung samples as described previously (3, 37). Similarly, the finding of elevated collagen levels through biochemical analysis parallels earlier studies showing higher collagen content through digital image analysis (3). Increased collagen deposition during the fibrotic response is a recognized indicator of disease progression (38). In concert, the release of bioactive collagen fragments that are revealed during the synthesis, deposition and turnover of ECM molecules, measured in the circulation or bronchoalveolar lavage, have also been recognized as potential biomarkers of disease activity in fibrotic lung diseases including in IPF (39-41) and ARDS (42-44). However, these studies only considered the total amount of collagen in study endpoints, rather than detailing the characteristics of the collagen fibers and how the collagen network is assembled and interlinked within the tissue.

Collagen fiber characteristics and the organization of the fiber network within lung tissue can be described through analysis of high magnification microscopy images. Properties such as the individual fiber length, number of fiber end points or curviness of the fibers, and more global fiber network parameters including the heterogeneity of fiber network patterns and the density of the matrix build a picture of the organization and assembly of the collagen in the lung tissue. The shape and complexity of this collagen fiber network is important for regulating cell responses (45). For example, the shape of collagen fibers can influence macrophage morphology and migration; with globular collagen fibers supporting more filipodia but a fibrous network inducing actin-rich protrusions and increased transmigration (46). Similarly, the topography of the fiber network influences mesenchymal cell fibrotic responses (19, 29, 47). Our digital image analysis revealed that several parameters associated with collagen fiber organization (collagen fiber length, branch points, end points and curvature, structural network heterogeneity and degree of high-density matrix) were increased in NR-ARDS and IPF lung tissues, but to different degrees. These observed differences in the collagen network assembly between NR-ARDS and IPF provide greater insight to the pathological environment than the increase in the absolute amount of collagen present in these lungs. These observations have implications for understanding of the similarities and differences in NR-ARDS and IPF fibrotic lung microenvironments, potentially elucidating disease perpetuating mechanisms.

The formation of the collagen fiber network is driven by complex processes. It is regulated by fiber proximity, spatial arrangement of supporting smaller matrix associated proteins and crosslinking of collagen fibers. In the extracellular space self-assembly of tropocollagen molecules (following cleavage of the pro-collagen pro-peptides) forms fibrils, followed by enzymatic crosslinking of lysine and hydroxylysine residues by the LOX family of enzymes to form fibers. Reactive aldehydes, that are the result of the LOX crosslinking reaction, then crosslink with aldehydes on adjacent collagen fibers that further condense to form a mature trivalent crosslink (**Figure 4A**) (48). This condensing process occurs over weeks-months (49). The most common of these mature trivalent crosslinks encompass hydroxylysyl pyridinoline and lysyl pyridinoline crosslinks. This is dependent on appropriate organization of the collagen fibrils before LOX activation for crosslinks to be formed, indicating an interrelationship between collagen crosslinking and structural organization (49). Stable mature collagen crosslinks increase the stiffness of the collagen network and make it more resistant to proteolytic cleavage and therefore less reversible. In our study we observed the highest number of mature crosslinks per collagen molecule in IPF tissues, but these were also present to a lesser extent in NR-ARDS. Interestingly, there was a correlation between the amount of lung collagen and the level of crosslinks per collagen molecule in IPF, but an inverse relationship was present in NR-ARDS, possibly reflecting the shorter time in which the trivalent crosslinks had to condense in NR-ARDS.

The LOX and lysyl hydroxylase enzyme families are essential for initiation of the process underlying collagen crosslinking. While defined roles have not been assigned to the different family members, LOXL2 has been shown to increase mature pyridinoline crosslink formation in bovine menisci (48). Pharmacological inhibition of the LOX family or LOXL2 specifically reduces collagen network fiber formation, matrix/tissue stiffening and improves lung function in model systems (29, 32). While inhibition of LOXL2 with simtuzumab was not effective in IPF patients (50), where the mature collagen crosslinks are well established such an approach may have potential value in NR-ARDS or earlier in the lung fibrosis disease courses.

Collectively, our results illustrate how the ECM structure varies between lung diseases associated with the presence of fibrosis. The altered collagen organization and (lack of) mature collagen crosslinking in both NC- and C-ARDS separate them from diseases with irreversible fibrotic changes, such as IPF. Nevertheless, this study has limitations. The patient-derived samples from different groups we had access to are not all derived from explanted lungs after lung transplantation, as most NC-ARDS group samples were derived post-mortem. As biological death is associated with many changes in the tissues and cells, this potentially alters ECM organization. Image-based digital analysis, albeit a strong tool, does not provide mechanistic insights, which is a weakness of this study.

Prolonged and NR-ARDS, although ill-defined (8), is accompanied with extensive tissue remodeling and collagen deposition (3). When ARDS does not resolve, despite treatment of the underlying cause, fibrotic changes on HRCT can lead to the assumption of irreversibility, prompting withdrawal of supportive treatment or escalating care towards lung transplantation. Our data might not support this implied irreversibility. Limiting tissue remodeling through antifibrotic treatment in an earlier phase might be of potential benefit in these patients to improve outcome (51).

## Conclusion

In summary, we examined how the collagen structural organization and crosslinking differed in lung tissues from patients with IPF and NR-ARDS and compared these with control counterparts. We revealed how collagen organization differed despite commonly high collagen levels in the diseased tissues and that fibrotic appearing regions in these diseases do not share a common makeup regarding collagen crosslinking maturity. Considering the limited number of therapeutic options for IPF and NR-ARDS, these results highlight important differences between these conditions and represent a step forward in determining the irreversibility of end-stage disease, as well as in identifying potential directions for specialized treatment strategies to reverse the fibrotic response.

## Acknowledgments

Authors thank Mr. Albano Tosato for assistance in preparation of figures.

## Data availability

The raw data used to generate the findings of this study are available from the corresponding authors upon reasonable request.

## Author Contributions

J.P. and J.K.B. conceived and designed the study. M.N., O.A.C., T.B., R.M.J., M.C.M., and R.H. performed experiments and/or acquired data. P.J.W. and J.P. acquired clinical data and/or contributed essential patient-derived materials. M.N., O.A.C., T.B., J.M.V., J.P., and J.K.B. analyzed and interpreted the data. M.N. drafted the manuscript and prepared the figures. M.N., O.A.C., T.B., J.M.V., R.M.J., M.C.M., R.H., P.J.W., J.P., and J.K.B. critically revised the manuscript for important intellectual content. J.P. and J.K.B. supervised the study and obtained funding. All authors approved the final version of the manuscript and agree to be accountable for all aspects of the work.

