## Supplementary Figures 1-6 for "Collagen crosslinking and organizational patterns reflect common disease processes in idiopathic pulmonary fibrosis and non-resolving acute respiratory distress syndrome"

### Supplementary material:

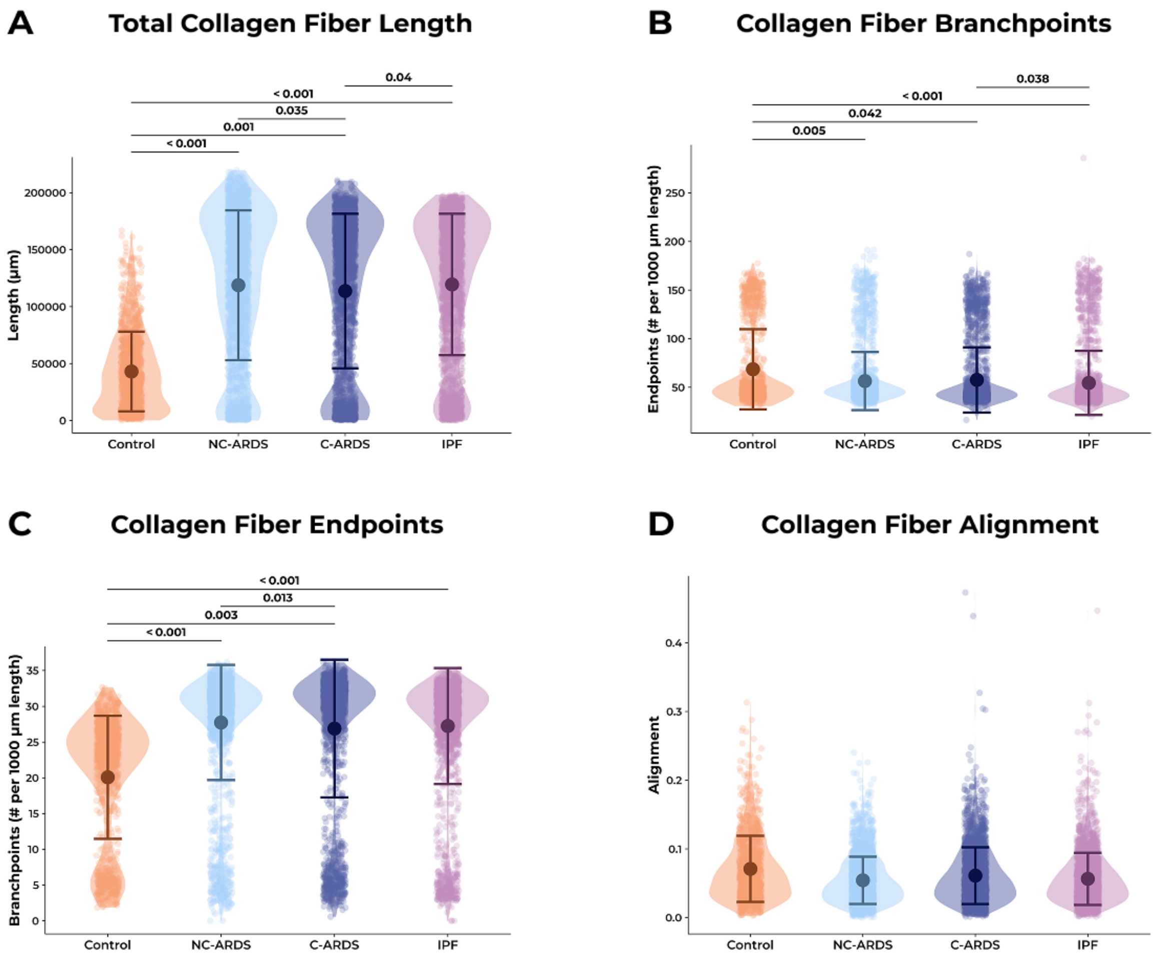


***Supplementary Figure 1:*** *Altered collagen assembly fiber metrics differed in ARDS and IPF lungs. The collagen organization analysis was performed through TWOMBLI analysis on images of PSR-stained lung sections. Quantification of A) total collagen fiber length, B) collagen fiber branchpoints, C) collagen fiber endpoints, D) collagen fiber alignment. Each dot with a smaller diameter represents one biological replicate. Data show mean ± SD. The analysis included 6 control individuals, 10 patients with NC-ARDS, 9 patients with C-ARDS, and 9 patients with IPF. Total image tiles analyzed, with per-individual ranges in parentheses, were as follows: control, 1,087 (90 to 415); NC-ARDS, 2,406 (132 to 405); C-ARDS, 2,374 (102 to 552); and IPF, 2,349 (101 to 372). A mixed model analysis was used to assess the statistical significances while accounting the variance from multiple images per sample. ARDS: Acute respiratory distress syndrome, C-ARDS: SARS-CoV-2 disease (COVID)-related ARDS, IPF: Idiopathic pulmonary fibrosis, NC-ARDS: non-COVID-related ARDS.*

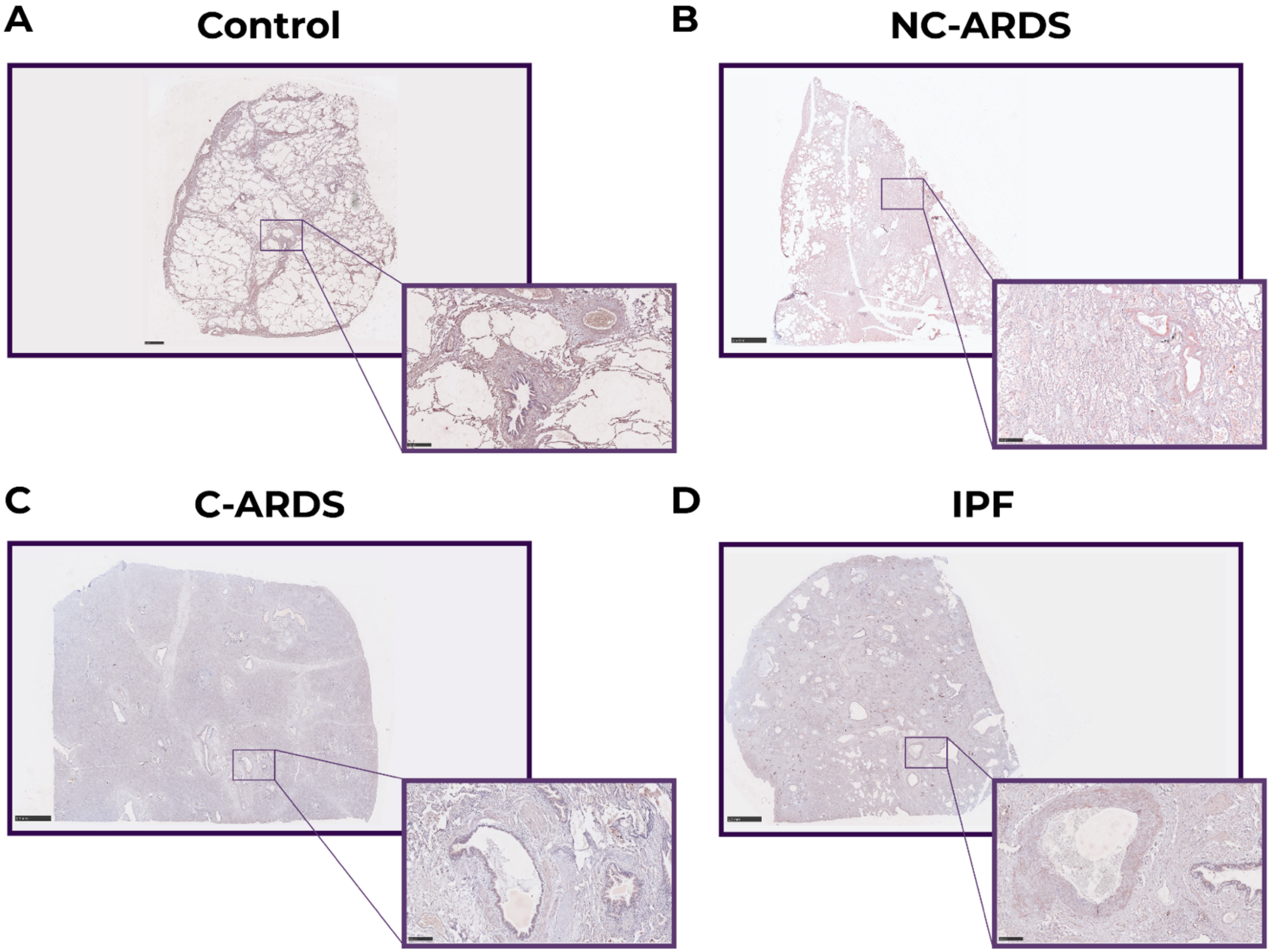


***Supplementary Figure 2****: LOX expression in lung tissues. Representative images of immunohistochemical staining against LOX, shown in brown, on A) control, B) NC-ARDS, C) C-ARDS, and D) IPF lung tissue. Scale bars: 2.5 mm for zoomed out panels and 150 µm for zoomed in panels. ARDS: Acute respiratory distress syndrome, C-ARDS: SARS-CoV-2 disease (COVID)-related ARDS, IPF: Idiopathic pulmonary fibrosis. LOX: Lysyl oxidase. LOXL: LOX-like, NC-ARDS: non-COVID-related ARDS.*

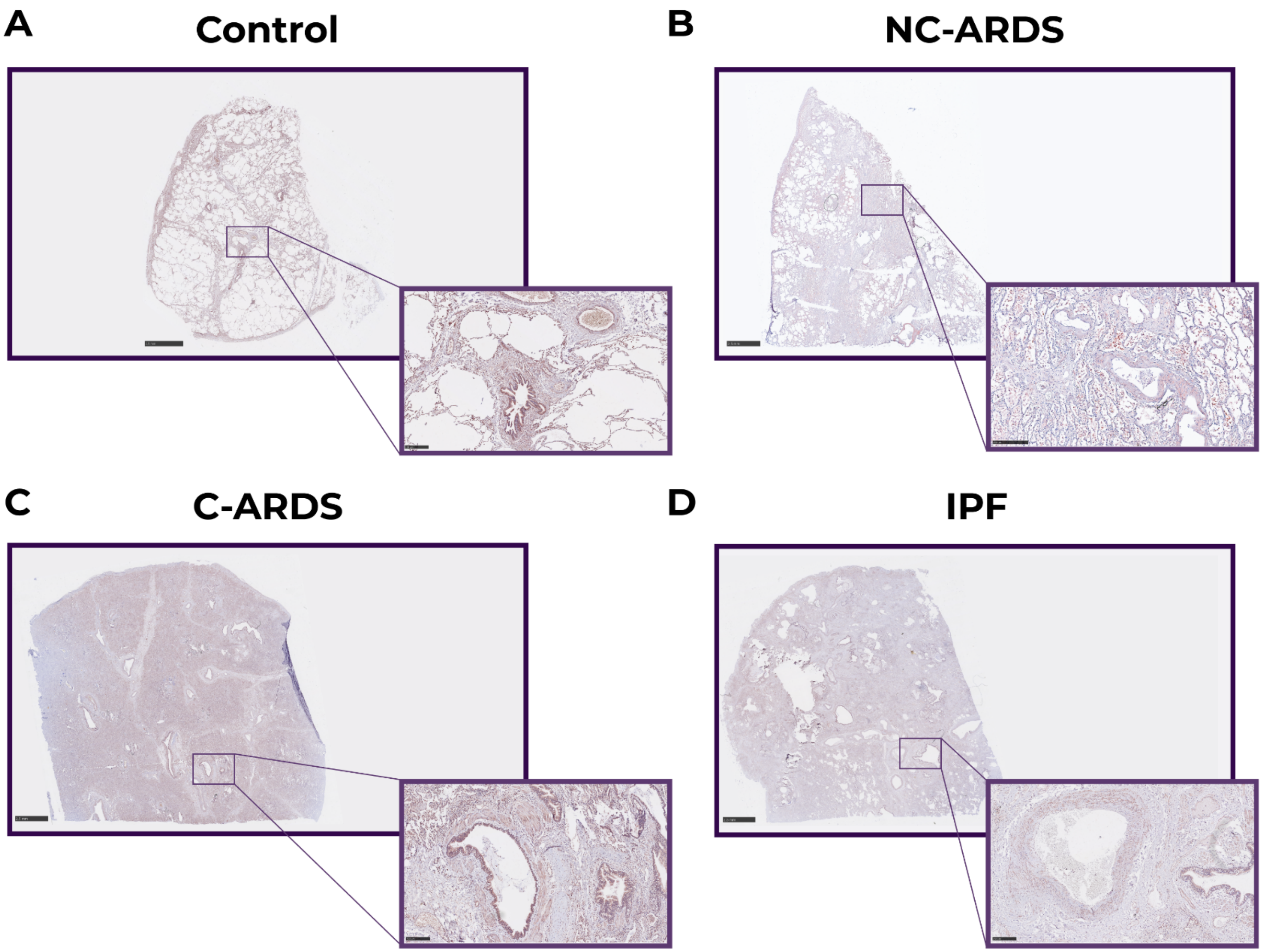


***Supplementary Figure 3***: *LOXL-1 expression in lung tissues. Representative images of immunohistochemical staining against LOXL-1, shown in brown, on A) control, B) NC-ARDS, C) C-ARDS, and D) IPF lung tissue. Scale bars: 2.5 mm for zoomed out panels and 150 µm for zoomed in panels. ARDS: Acute respiratory distress syndrome, C-ARDS: SARS-CoV-2 disease (COVID)-related ARDS, IPF: Idiopathic pulmonary fibrosis. LOX: Lysyl oxidase. LOXL: LOX-like, NC-ARDS: non-COVID-related ARDS.*

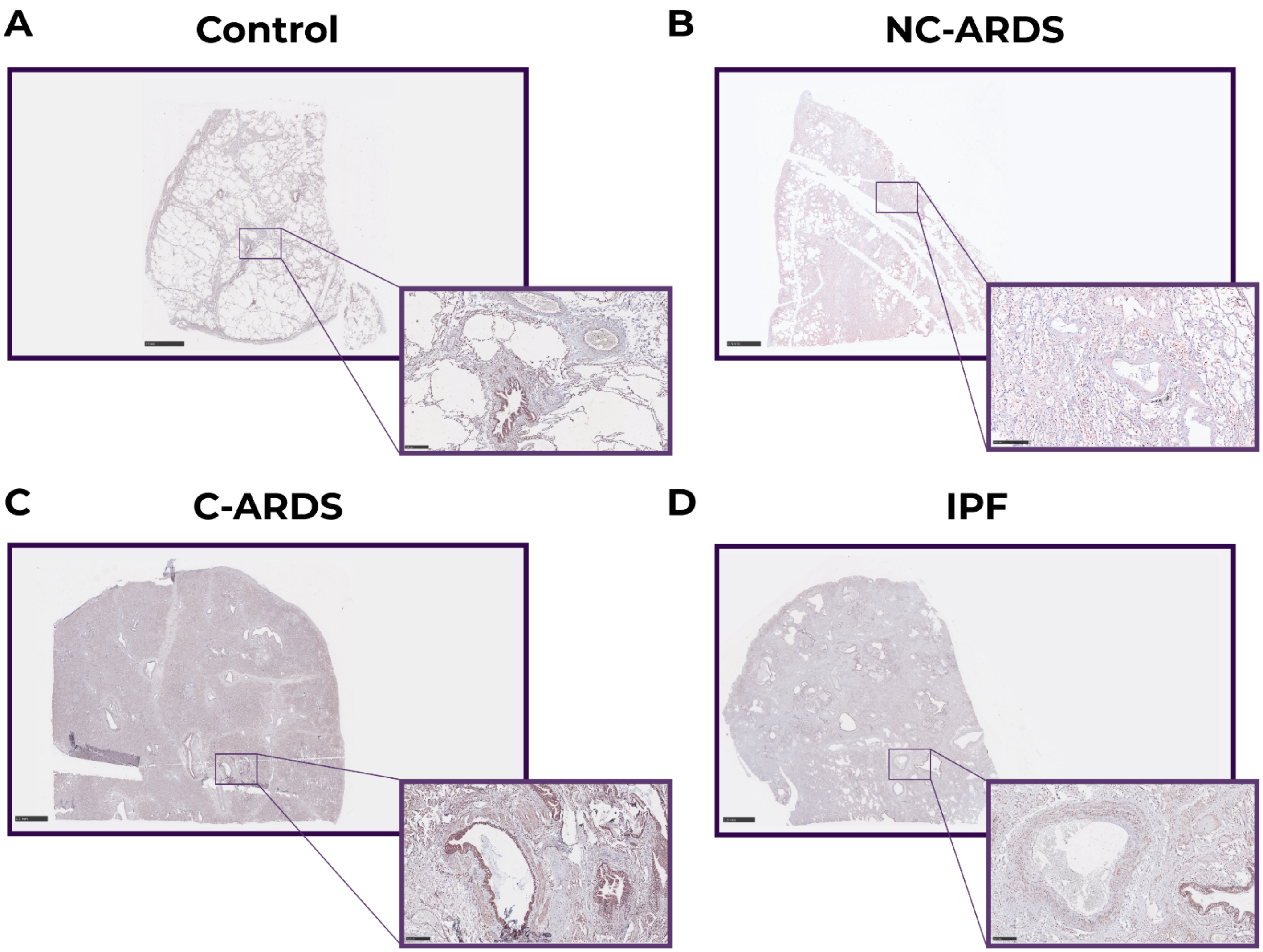


***Supplementary Figure 4***: *LOXL-2 expression in lung tissues. Representative images of immunohistochemical staining against LOXL-1, shown in brown, on A) control, B) NC-ARDS, C) C-ARDS, and D) IPF lung tissue. Scale bars: 2.5 mm for zoomed out panels and 150 µm for zoomed in panels. ARDS: Acute respiratory distress syndrome, C-ARDS: SARS-CoV-2 disease (COVID)-related ARDS, IPF: Idiopathic pulmonary fibrosis. LOX: Lysyl oxidase. LOXL: LOX-like, NC-ARDS: non-COVID-related ARDS.*

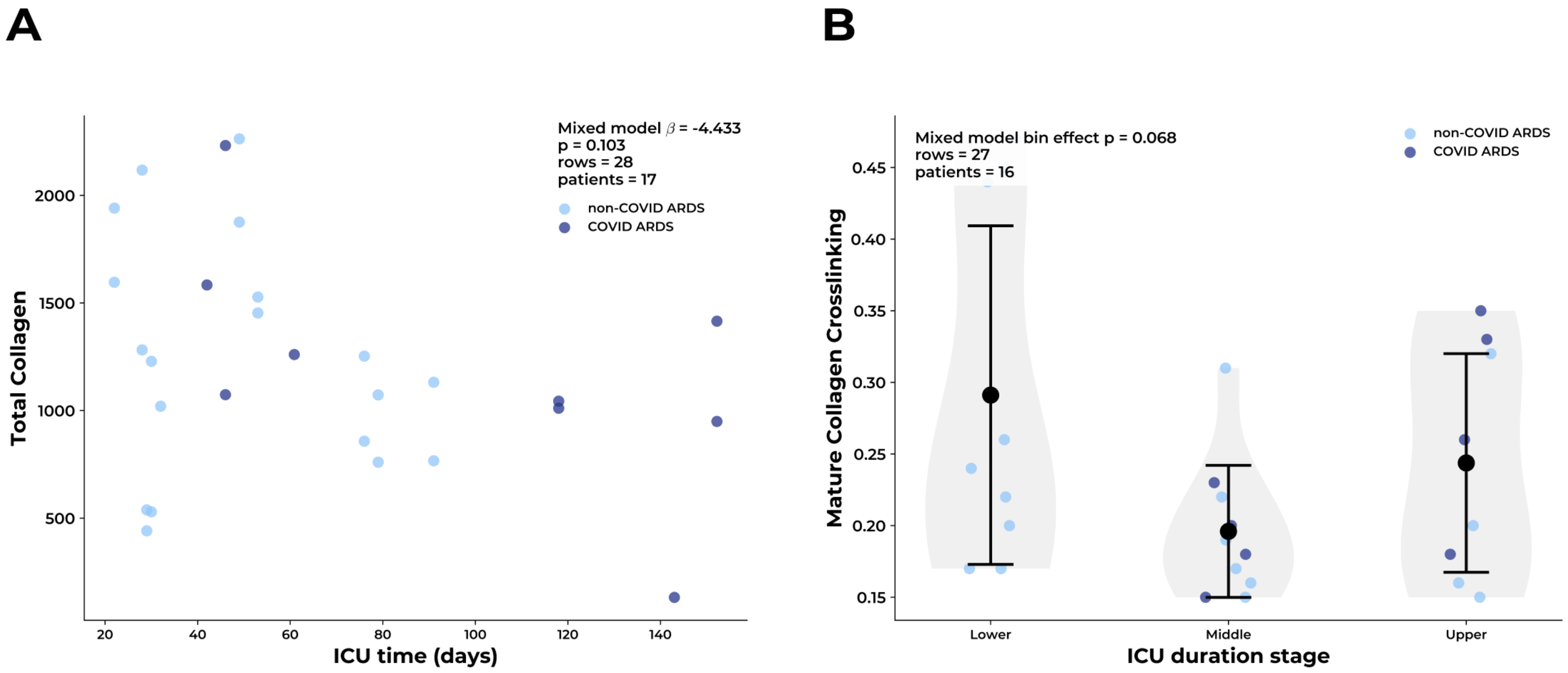


***Supplementary Figure 5:*** *Relationships among collagen remodeling metrics and ICU duration. A) Association between ICU duration and total collagen in ARDS donors, separated by COVID and non-COVID ARDS. C) Mature collagen crosslinking across ICU duration stages in ARDS donors using data-driven tertile cut-points: Lower: ≤38.67; Middle: 38.67-76.00; and Upper: >76.00 days. Individual points represent individual donors, in which color indicates ARDS subgroup. In panel C, data are represented as mean ± SD. Sample sizes for panel B were n=7 for control, n=19 for non-COVID ARDS, n=11 for COVID ARDS, and n=10 for IPF, derived from 6 control individuals, 10 patients with non-COVID ARDS, 8 patients with COVID ARDS, and 10 patients with IPF. Sample sizes for panel C were n=19 for non-COVID ARDS and n=8 for COVID ARDS, derived from 10 patients with non-COVID ARDS and 6 patients with COVID ARDS. ARDS: Acute respiratory distress syndrome.*

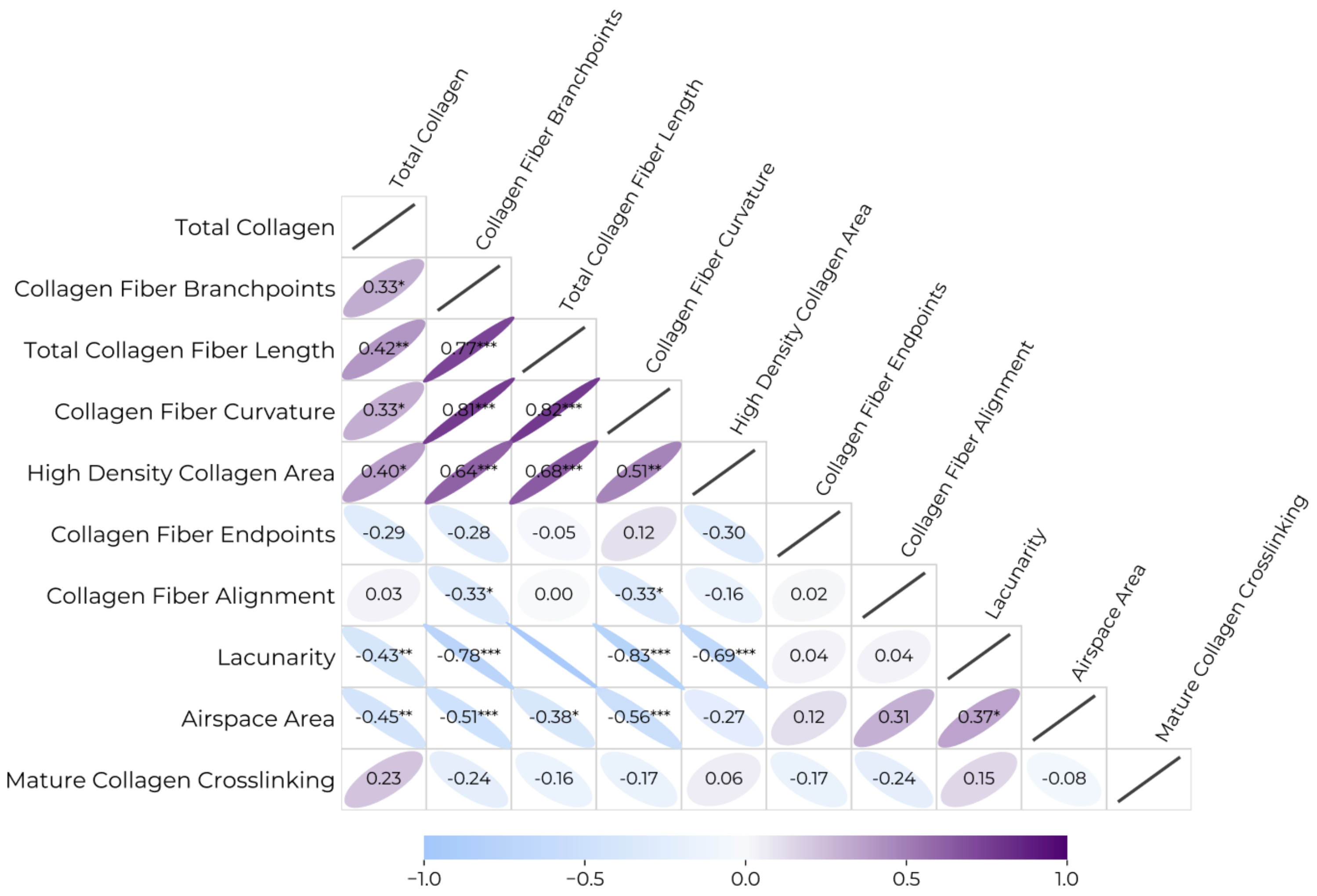


***Supplementary Figure 6:*** *Correlations among collagen structural and tissue architecture metrics used in principle component analysis (PCA). Correlation heatmap of collagen structural and tissue architecture metrics used in the PCA. Pairwise Spearman correlation coefficients (ρ) are shown for donor-level, median-aggregated features. Color indicates the direction and magnitude of the correlation: blue indicates negative correlation and purple indicates positive correlation. Numeric values correspond to correlation coefficients, and asterisks denote significance: *p<0.05, **p<0.01, ***p<0.001. Only the lower triangle is displayed for clarity.*
